# Crosslink Strength Governs Yielding Behavior in Dynamically Crosslinked Hydrogels

**DOI:** 10.1101/2024.08.16.608324

**Authors:** Noah Eckman, Abigail K. Grosskopf, Grace Jiang, Krutarth Kamani, Michelle S. Huang, Brigitte Schmittlein, Sarah C. Heilshorn, Simon Rogers, Eric A. Appel

## Abstract

Yielding of dynamically crosslinked hydrogels, or transition between a solid-like and liquid-like state, allows facile injection and utility in translational biomedical applications including delivery of therapeutic cells. Unfortunately, characterizing the time-varying nature of the transition has not been attempted, nor are there design rules for understanding the effects of yielding on encapsulated cells. Here, we unveil underlying molecular mechanisms governing the yielding transition of dynamically crosslinked gels currently being researched for use in cell therapy. We demonstrate through nonlinear rheological characterization that the network dynamics of the dynamic hydrogels dictate the speed and character of their yielding transition. Rheological testing of these materials reveals unexpected elastic strain stiffening during yielding, as well as characterizing the rapidity of the yielding transition. A slower yielding speed explains enhanced protection of directly injected cells from shear forces, highlighting the importance of mechanical characterization of all phases of yield-stress biomaterials.

**Significance Statement:** Many direct-injection cell therapies suffer from poor viability of injected cells due to membrane disruption by shear stresses during injection. Injectable hydrogels with dynamic crosslinks can increase viability during injection, utilizing the ability of these materials to undergo a reversible solid-to-liquid shear-rate dependent yielding transition. Much effort has been applied to understanding the yielding transition of these materials, yet design rules for connecting the character of yielding to viability of injected cells has not been established. Here, we apply rheological testing to show that faster network dynamics result in a more gradual yielding transition, affording greater protection to cells during injection. This work introduces a new rheological protocol for the development of injectable hydrogels for cell therapy.

## Introduction

Dynamically crosslinked hydrogels (DCHs) are water-swelled polymer networks which are not permanently crosslinked, but are instead formed by transient associations between polymer chains^1^. The increased interest in the unique capabilities of these materials have given rise to their use as diverse biomedical tools, having been used in the fields of 3D printing, wound repair, adoptive cell therapy, drug delivery, vaccines, and beyond^2–4^. Dynamic associations may allow these hydrogels to be injectable, respond to physiological cues, and allow for cellular mobility and nutrient transport while affording structural stability. The viscoelasticity and yield-stress behavior of these dynamic networks additionally afford protection to cells from shear stress-induced lysis during injection, improving efficacy of cell delivery technologies^5,6^. Unfortunately, the transient associations of the network imbue the material with complex rheological properties that have been historically challenging to characterize^7^. Theory-based approaches or fundamental rheological studies which analyze the mechanical properties of transiently crosslinked networks usually rely on intentionally architected materials or simplifications, often conceptualizing the network as a single-mode Maxwell model to fit stress relaxation and linear frequency sweeps and calculate kinetic parameters^8–11^. Yet, novel materials being implemented in biological studies rarely obey model behavior, and exhibit highly complex rheological features, so qualitative descriptions are necessary^12,13^. Additionally, lack of standardization in mechanical and cell viability testing makes it difficult to compare materials head-to-head.

In general, DCHs are elastoviscoplastic materials which also exhibit time-dependent (thixotropic) elastic and viscous behavior^14–16^. The yielding transition of these materials is of particular interest for their biomedical use case^7^. The transition of DCHs from a solid-like state to a liquid-like state corresponds to the breaking of the transient crosslinks. This transition is not instantaneous in time nor does it occur at a single well-defined yield stress value, creating complications in the characterization of the transition^17^. During injection, a material begins in a quiescent equilibrium state before being subjected to extremely high shear rates (>10^4^ s^-1^) during flow, breaking the transient associations. After injection, the crosslinks reform on a characteristic time scale, during which the material is subject to a creep (constant stress) condition as it is confined within a tissue or under the skin. Understanding the material response during all operating conditions is critical to evaluating a DCH, especially the yielding transition, but typically, material evaluation focuses on the linear (pre-yield) viscoelastic regime. Indeed, many authors only report a single value (G’, the storage modulus) for the mechanical characterization of DCH materials. Large-amplitude oscillatory shear (LAOS) is an attractive option for evaluation of a range of mechanical properties, as it probes the pre-yield, yielding, and post-yield behavior of a given material^18,19^. Unfortunately, physical interpretation of LAOS data, especially in the non-linear regime where the stress response becomes non-sinusoidal, is difficult^19,20^. Recently a suite of techniques has emerged, including Sequence of Physical Process (SPP) analysis^21,22^, which aid in the interpretation of LAOS data. Recent advances have also evaluated the yielding of soft materials on a spectrum from brittle to ductile, where brittle refers to a sudden yielding transition whereas ductile materials yield gradually^23^.

In this work, we applied SPP analysis of LAOS data to three clinically relevant DCHs with various crosslink chemistries. We use a whole-waveform approach to visualize stress-strain or time-dependent modulus and phase angle data which is usually discarded in LAOS experiments. SPP analysis enables us to avoid relying on interpretation of higher-order sinusoids, which can be difficult to understand experimentally. The networks we evaluated exhibit similar linear viscoelastic properties, including their storage modulus G’, which is proportional to crosslink density in dissociative polymer networks^24^. We characterized the yielding transition of the networks and show critical qualitative and quantitative differences in the yielding process explained by the crosslink thermodynamics. Additionally, we use a geometric technique for stress decomposition to examine the time-varying elastic responses of the material and show an elastically derived strain overshoot which corresponds to the activation energy needed to break crosslinks and begin flow. SPP analysis was used to calculate instantaneous phase angles and phase angle velocities, which we correlated to a yielding speed, or abruptness of the yielding transition^23^. To verify the finding of an increased yielding speed, or more fracture-like yielding in materials with stronger crosslinks, we tested the protection of encapsulated cells to injection at high shear rates and found that gels with more ductile yielding transitions afford greater protection of cells from shear forces (**Figure 1A**). By applying sophisticated rheological techniques to complex materials which have been put into practice in the biomaterials literature, this work serves to bridge the gap between rheological studies which examine model materials and the biomedical materials science community.

**Figure 1.**
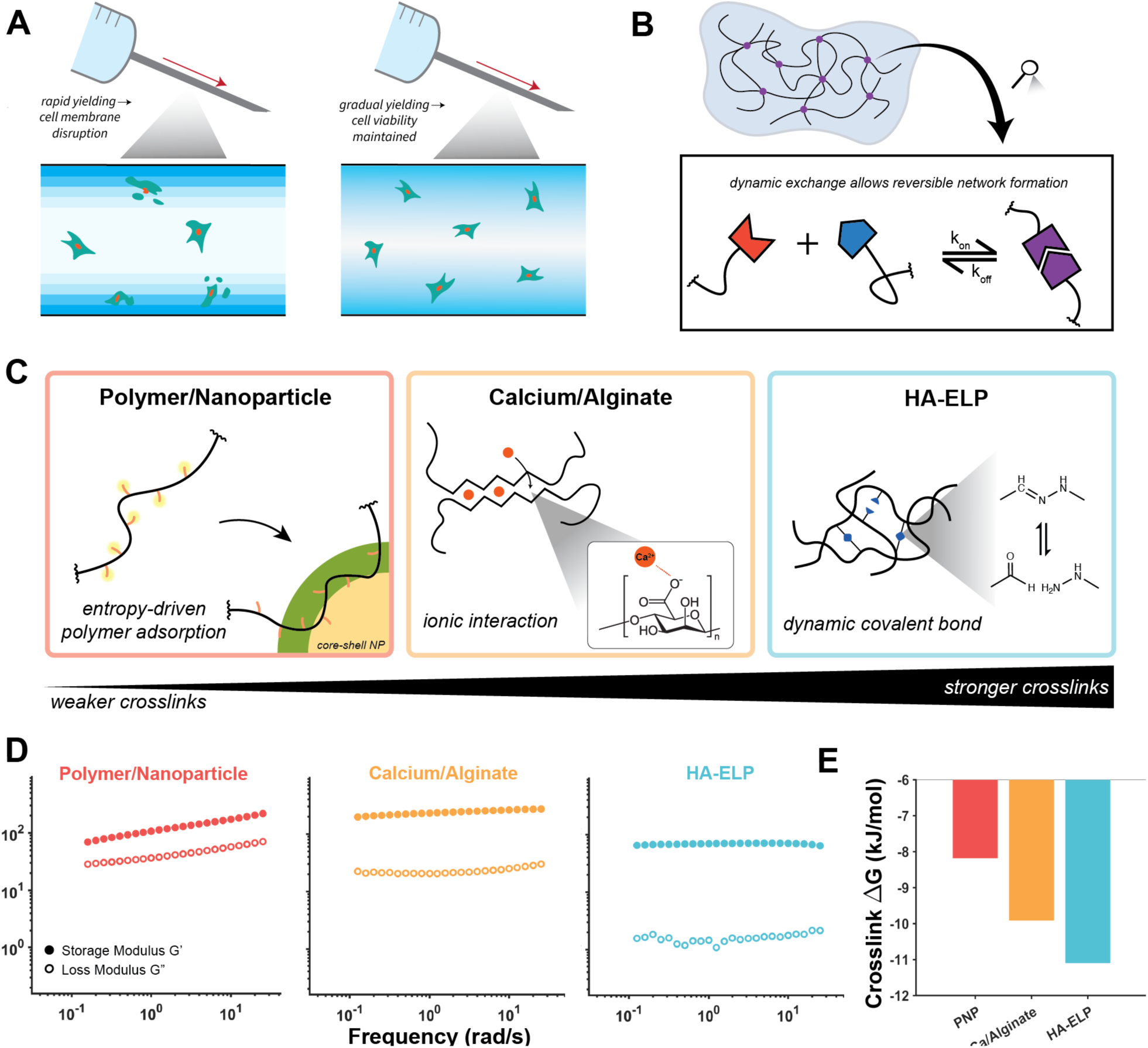
Linear rheological comparison of dynamically crosslinked hydrogels. (A) Hydrogel-encapsulated cell viability is impacted by the rapidity of yielding: in rapidly-yielding gels, velocity gradients disrupt cell membranes, while in gradually yielding gels, cell viability is maintained. (B) Dynamically crosslinked hydrogels (DCHs) are formed by transient crosslinks in which dynamic exchange allows reversible network formation. (C) Schematic of the three DCH highlighted in this investigation. Polymer/nanoparticle hydrogels are formed by entropy-driven adsorption of a hydrophobically-modified cellulose polymer onto the surface of core-shell (poly)ethylene glycol—(poly)lactic acid nanoparticles. Calcium/alginate hydrogels are formed by ionic interactions in an “eggbox” architecture between divalent cations and the anionic monomers. Hyaluronic acid—Elastin-like protein (HA-ELP) hydrogels are formed by hydrazone bond formation between the aldehyde-modified HA and hydrazine groups on ELP. (D) Frequency sweeps of the three DCH materials, showing similar linear viscoelastic gel-like plateau behavior. (E) Comparison of the crosslink equilibrium constant for the DCH materials, showing increasing crosslink strength of the materials, but all reversible. Data adapted from Refs. (8) and (26).

## Results and Discussion

### Comparison of hydrogel systems

To apply the SPP-LAOS approach to multiple relevant biomaterials platforms, we chose three separate DCHs which have distinct crosslinking mechanisms (**Figure 1B, C**). The three systems can be ordered from weakest crosslinks to strongest crosslinks, although all the crosslink chemistries have dynamic character. Polymer-nanoparticle (PNP) hydrogels are formed by a multivalent, noncovalent entropy-driven association between a hydrophobically-modified polymer which adsorbs to the surface of nanoparticles which act as crosslinkers^9^. Alginate hydrogels are formed through the dimerization of two polymer chains (conceptualized through the “egg-box” model) by ionic interactions between Ca^2+^ ions and anionic monomers along the polysaccharide backbone^25^. Finally, hyaluronic acid/elastin-like protein (HA-ELP) hydrogels are formed by crosslinking of hyaluronic acid (HA) polymers by dynamic hydrazone bonds formed between aldehyde modification groups on HA and hydrazine modification groups on ELP^26,27^. The equilibrium constants (K_eq_) of the three systems range between 10^3^ and 10^5^, representing crosslinks which are stable enough to form percolated gel networks, but which can be broken upon application of mechanical stresses relevant for processes such as injection. The linear rheological properties of the three networks are all similar (**Figure 1D**), with relatively flat frequency responses of both dynamic moduli along the experimentally relevant frequency range. These materials do not exhibit time-temperature superposition, so collection of additional frequency data is not feasible^9^. The slope of the storage modulus decreases from low to high crosslink strength, which could be attributed to either more arrested behavior or decreased dispersity of the polymer networks between the hydroxypropylmethylcellulose (HPMC) polymer (highly disperse) and ELP (monodisperse). Interestingly, even though the crosslinking strategies of the networks are quite different and correspondingly have a range of crosslink free energies of binding (**Figure 1E**), all three have an approximately similar number crosslink densities and linear elastic modulus *G*’_*p*_ values in accordance with network theory (Supplemental Section 1). From low to high crosslink strength, the loss modulus *G*” decreases, which could be attributed to stronger confinement effects on solvent molecules. Crucially, the linear characterization of the materials fails to describe their yielding transition and flow properties, so additional characterization was required.

### LAOS elucidation of material regimes

Because the implementation of these injectable materials (e.g., by injection) requires control over their behavior before, during, and after yielding, we sought to implement a characterization technique which could probe multiple regimes. We used large-amplitude oscillatory shear to characterize the different material regimes of the DCHs. The measurement was taken by performing oscillatory rheology sweeps from small to large amplitudes at low frequency to avoid inertial artifacts. 1-dimensional amplitude sweeps are shown in **Figure 2A**, with the dynamic moduli *G*’ and *G*” remaining flat in the linear regime and then falling off in the nonlinear regime as the gel structure breaks down during yielding. Yet, in the nonlinear viscoelastic regime, the moduli lose their significance as in-phase and out-of-phase portions of the stress response. Therefore, we used parametric Lissajous stress/strain curves to examine the material behavior regimes (**Figure 2B**). In the elastic regime, the Lissajous curves approximate straight lines with slope *G*’, since *tan*(*δ*) < 1. As the material yields, the curves enclose more area as the viscous response grows in importance, and non-linear effects emerge. Finally, in the flow regime, successive curves have a constant value of stress for many values of strain, seen by the curves flattening at the top; this may represent evidence of shear-banding^28^. In **Figure 2C**, we added the strain rate (simply the time derivative of the strain) as a third spatial variable to better illustrate the yielding transition. The first striking difference between the curves is the presence of a stress overshoot in some formulations evaluated. PNP gels didn’t exhibit a stress overshoot, as the curves increased in strain and approached a plateau value for stress. In contrast, the Ca/Alginate gels exhibited a robust stress overshoot, as the elastic region curves attained a higher value of stress than the flow plateau by about 2-fold. Finally, the HA-ELP gels exhibited an extremely large overshoot value, obtaining a stress in the elastic regime nearly 10 times higher than in the terminal flow regime. The yielding transition can also be qualitatively assessed from these 3D curves. We conceptualized the pre-yielding curves as lying on a shared plane then transitioning to an orthogonal plane once yielded (Supplemental Figure 4). From low to high crosslink strength, the materials exhibited a more gradual to more abrupt yielding transition. While it is difficult to pinpoint an exact strain for which the PNP gel yields, the Ca/Alginate and HA-ELP gels exhibited a more sudden transition, occurring largely between two successive amplitude strains. A sudden yielding transition may afford more precision and predictability in processing, whereas a gradual yielding transition may afford the advantage of less abrupt changes in the environment of encapsulated cells. These results show how visualization of amplitude sweeps in multiple dimensions can elucidate qualitative differences in the yielding transition of these materials.

**Figure 2.**
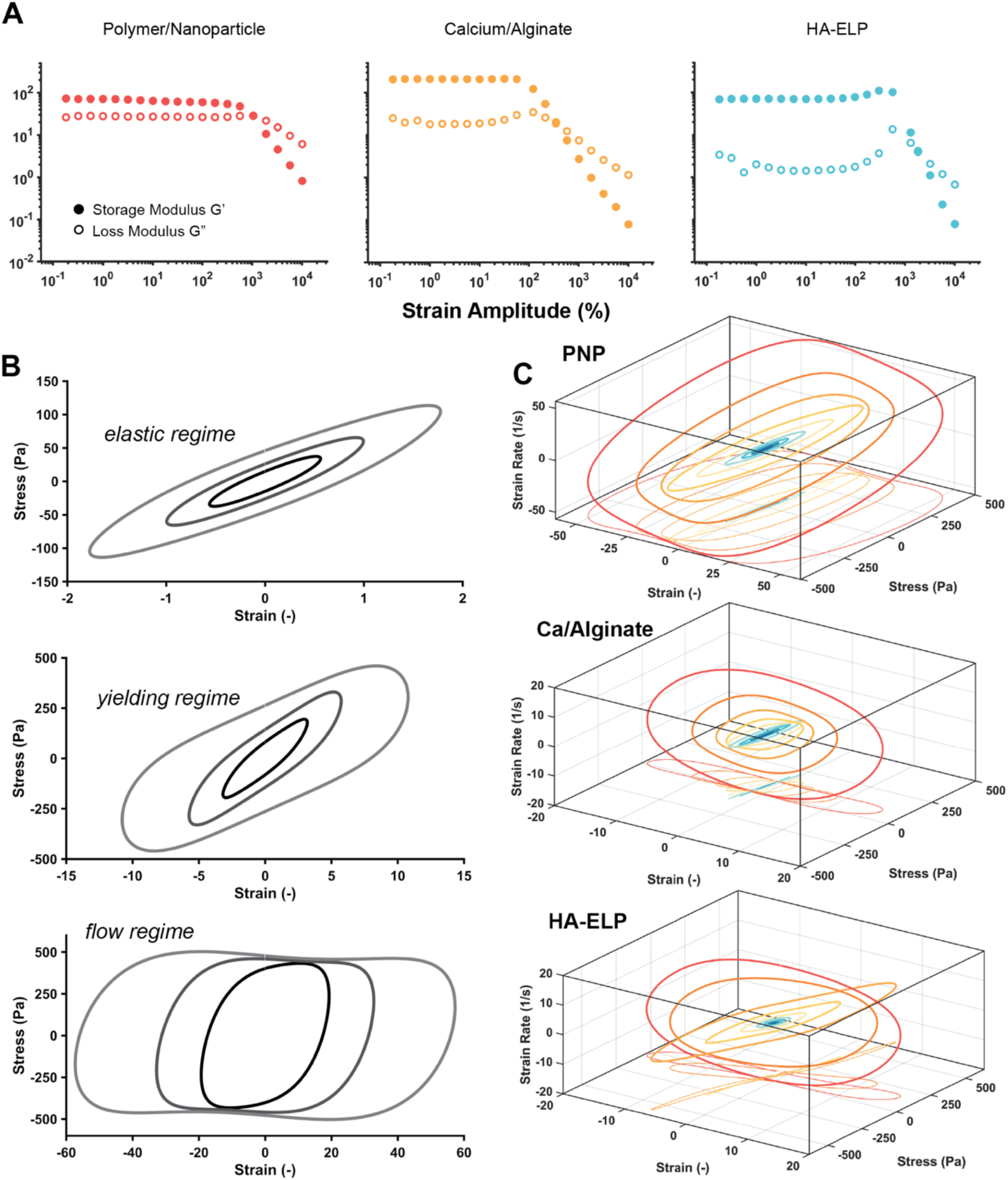
Large-amplitude oscillatory shear rheology. (A) Amplitude sweeps show the extent of the linear viscoelastic regime and subsequent yielding of the DCH materials at high strain. Beyond the linear viscoelastic regime, the dynamic moduli are fit to the fundamental mode of the strain oscillation and are not representative of the complete material response. (B) Illustration schematic of Lissajous curves which demonstrate the yielding transition from a mostly elastic material response to a mostly viscous material response. (C) 3-dimensional Lissajous curves for the DCH materials, illustrating the stress overshoot before yielding and which illustrate the gradual nature of yielding for weakly crosslinked materials, to the sudden yielding transition for strongly crosslinked materials.

### Elastic strain decomposition

Next, we used strain decomposition to better understand the physical origins of the observed stress overshoots. The elastic and viscous portions of the oscillation stress were calculated as follows^29^:

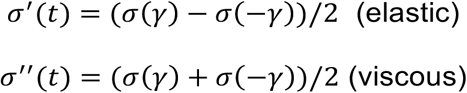

From there, we visualized the Lissajous curves for the time-varying elastic stress with respect to strain (**Figure 3A**). The curves transition from straight lines in the linear regime to exhibiting strongly non-linear effects at large *γ*_0_. For all the curves, we observed an initial increase in the maximum stress value before the material yields and the maximum stress declines to a plateau value. The magnitude of the elastic stress overshoot greatly increased between the three materials corresponding with increasing crosslink strength. We hypothesized the physical origin of the elastic stress overshoot in **Figure 3B**. At equilibrium, polymer strands are connected by the dynamic crosslinks. As strain increases, the strands are pulled apart, with increasing stress, until at a critical stress the crosslinks break, and the material begins to flow. Stronger crosslinks require a greater activation energy to break, and thus more stress is required to initiate flow. After the crosslinks have been broken, the residual elasticity is due to the intrinsic elasticity of the polymer chains. We hypothesize that the drastic reduction in the stress post-yield we observed in these materials is due to the onset of shear-banding^28^. For more strongly crosslinked gels, the onset of shear banding is more pronounced and results in a drop-off in stress as heterogeneities develop in the flow field.

**Figure 3.**
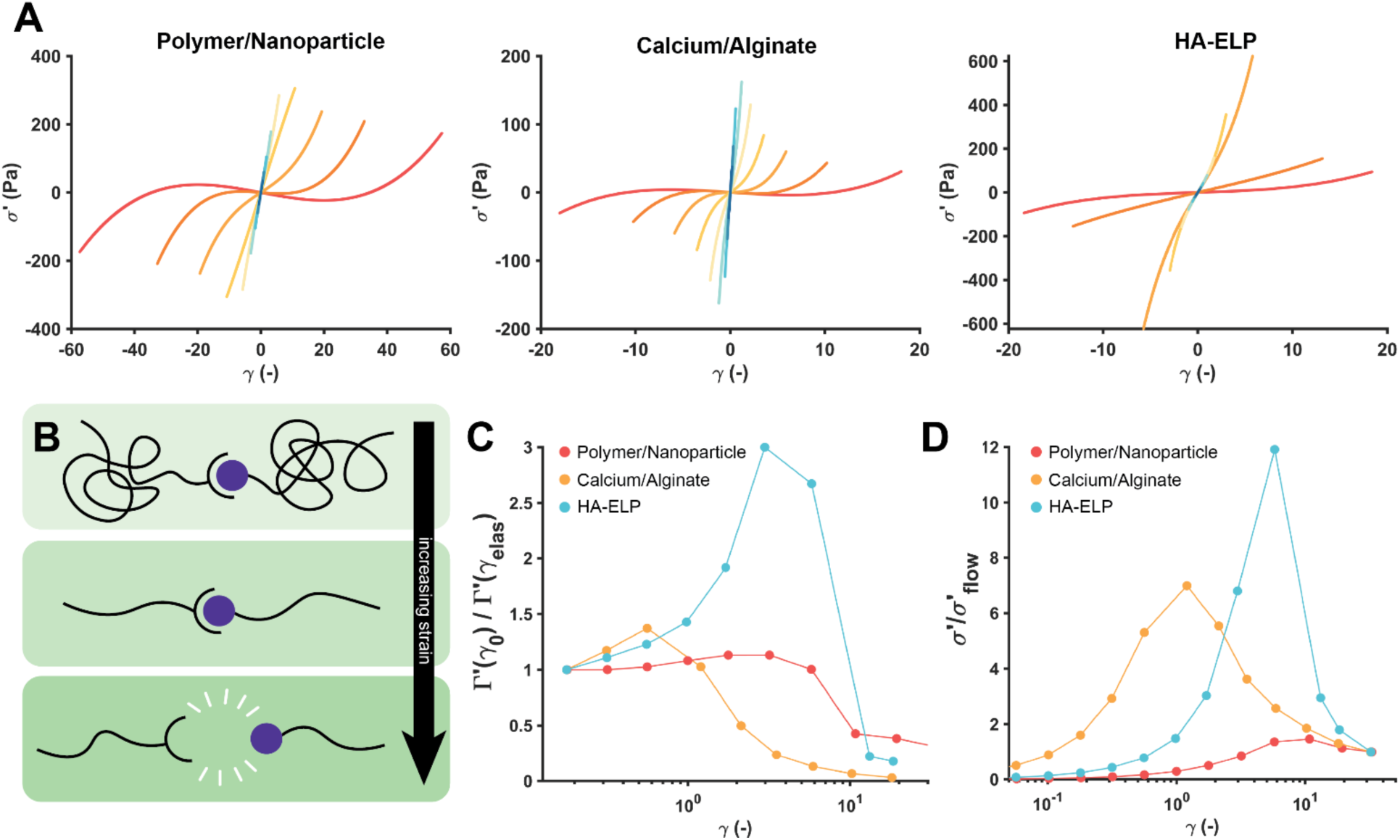
Molecular theory of the elastic stress overshoot. (A) Elastic stress decomposition from LAOS experiments. The overall stress response is decomposed into elastic and viscous portions, and here is plotted the Lissajous curve of the elastic stress against the strain for a series of oscillations. (B) Schematic of the molecular origin of the elastic stress overshoot. As extension of chains increases, the dynamic crosslinks continue to connect individual chains. At a critical strain, the crosslinks break allowing the network to flow. For weakly crosslinked materials, the crosslinks break at a lower chain extension. At high crosslink strength, the chains extend to a greater extent, causing a nonlinear increase in the stiffness before yielding. (C) Strain stiffening during yielding. Here, we compare the generalized elastic modulus evaluated at the maximum extension of each cycle, Γ′(*γ*_0_), to the stiffness in the linear viscoelastic regime Γ′(*γ*_*elas*_), which is equivalent to G’. Weakly crosslinked materials show little strain stiffening, while the strongest crosslinked materials undergo drastic pre-yield elastic strain stiffening. (D) Elastic stress overshoot. We compare the maximum value of the elastic stress for each oscillation to the terminal stress, approximately the yield stress of the material. Stronger crosslinks result in an overshoot above the terminal stress, caused by the increase in stiffness at high extension.

To better quantify the elastic stress overshoot, we adopted the definition of a generalized elastic modulus Γ′(*t*) = *σ*′/*γ*, measuring the ability of the material to store energy in the nonlinear regime. This definition eliminates the need to rely on a single harmonic fit to the stress, as is normally done for G′ and G″. We then compared Γ′ measured at *γ*_0_ (corresponding to maximum extension of elastic elements) to Γ′ at small angles, which is equivalent to G′ (**Figure 3C**). This approach measures the nonlinearly increased stiffness of the material at its most extended state, showing that the more strongly crosslinked gels undergo an elastic strain stiffening before a rapid decrease in stiffness post-yielding^30^. Importantly, we analyzed stress data from successive amplitudes, rather than identifying strain stiffening in one period^30^. Finally, in **Figure 3D**, we measured the ratio of the maximum of the elastic stress for a given amplitude to the value of elastic stress during flow (defined as the plateau value on the stress/strain Lissajous curves at post-yield *γ*_0_). This metric measures the effective elastic stress overshoot and demonstrates a trend from weakly crosslinked gels which display little stress overshoot, to more strongly crosslinked gels which have a stress overshoot of over 10 times the flow value. We hypothesized that this value describes the yielding behavior on an axis from “smooth” to “fracture-like”. Considering the extreme case for each type of material helps to explain this distinction. On one extreme, a liquid displays no yield stress as increasing strain acquires higher and higher values of stress. For an elastic solid, once the material fractures, the retained stress drops to zero. This spectrum of yielding behaviors highlights the difficulties in defining single yield stress values for dynamically associating networks.

It is also important to highlight the distinction between this elastic stress overshoot effect and the previously reported weak strain overshoot in the loss modulus during yielding, a common feature of rubbers and other soft materials. The viscous (G”) strain overshoot arises from the “de-caging” of solvent molecules as the structure of the network begins to break down and local yielding occurs, whereas we hypothesize the elastic strain stiffening effect to arise from chain extension in the pre-yielding regime before crosslinks are broken^31^. To investigate whether these effects are correlated, we tested the yielding behavior of other hydrogels that exhibit a weak strain overshoot in the loss modulus during yielding, but indeed which exhibit no elastic strain overshoot, thus implying the distinct and uncorrelated nature of these phenomena (**Supplemental Figure 3**). In the DCH materials we tested, the increasing crosslink strength also corresponds to greater viscous overshoot along with the elastic overshoot, but these trends are evidently not universal.

### Phase angle velocity and yielding speed

To further examine the rapidity of the yielding transition, we used time-varying Cole/Cole (G’/G”) plots to visualize the value of the dynamic modulus during each oscillation. Traditionally, the dynamic moduli are given in the nonlinear regime as fits to the fundamental mode of the stress, but this neglects non-linear effects. We took advantage of the SPP definition of the time-varying dynamic moduli *G_t_*′ and *G_t_*″, which correspond to the instantaneous material behavior in the non-linear regime and accounted for changes over the course of one oscillation, including yielding and un-yielding. Over the course of a half-oscillation, the material sweeps through one loop of the Lissajous curve as the instantaneous dynamic moduli *G_t_*′ and *G_t_*″ change with respect to time (**Figure 4A**). The instantaneous phase angle *δ*_*t*_ is defined by the orientation of the vector *G*^∗^ = *G*′ + *iG*′′ in the complex plane, with the phase angle velocity 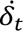 defined as the time derivative of *δ*_*t*_. In the linear viscoelastic limit, the time-varying moduli are constant, as the stress response is sinusoidal. As the material begins the yielding transition and nonlinear effects are introduced, the curves transition to a more liquid-dominated behavior (**Figure 4B**). We then defined the material yielding speed in the LAOS experiment as the maximum of the phase angle velocity during a single oscillation. **Figure 4C** shows the maximum yielding speed over the course of the LAOS experiment, illustrating the increased rapidity of the yielding transition with increasing crosslink strength and thereby corroborating the observations in **Figure 2C**.

**Figure 4.**
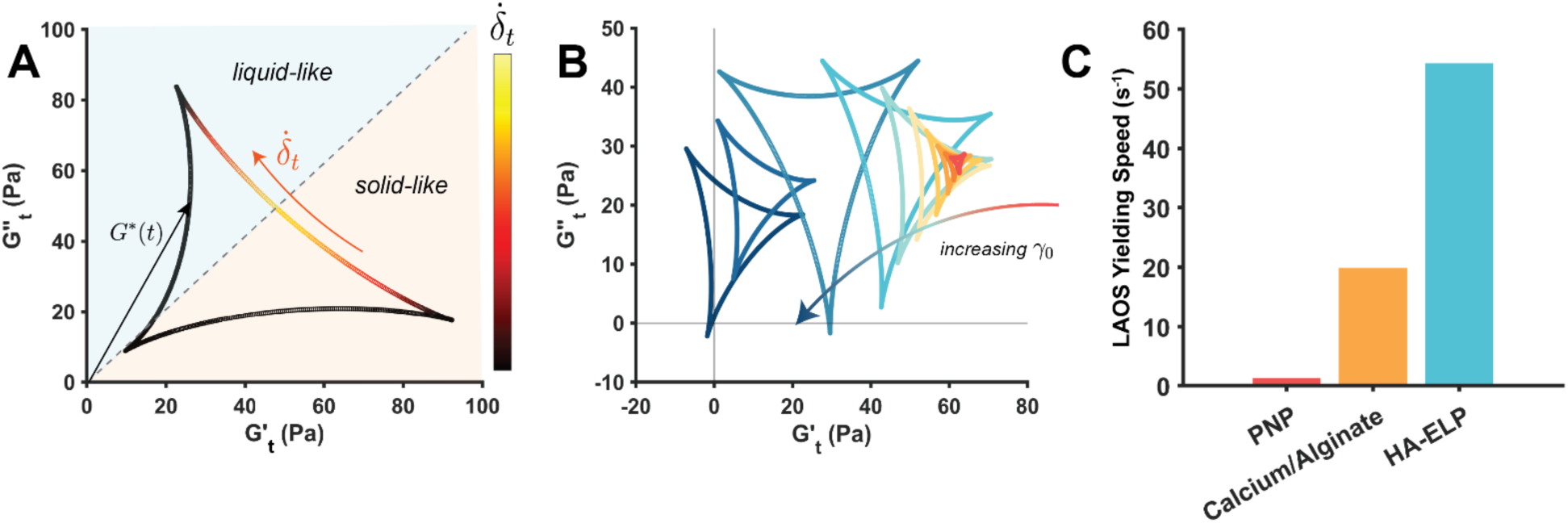
Speed of yielding as measured by Cole/Cole plots. (A) Schematic of a parametric Cole/Cole plot, which illustrates yielding within one oscillation. In the nonlinear regime, we calculate the instantaneous storage and loss moduli *G_t_*′ and *G*′′_*t*_. In this plot, the material starts out solid-like, with *G_t_*′ > *G*′′_*t*_. As the material yields, the curve sweeps counter-clockwise, crossing the line of tan(*δ*) = 1 as the response becomes liquid-like. The instantaneous speed of the phase angle, 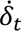, is shown here by the color of the curve at each point. (B) Overlay of Lissajous curves for an amplitude sweep, with cooler colors representing greater strain amplitude. In linear viscoelasticity, the curve is a single point, as the moduli are constant. As yielding proceeds, the curves transition to the left as the material becomes more liquid-like than solid-like. (C) Speed of yielding as calculated by the phase angle rate. We choose the Cole/Cole Lissajous curve which has the largest area and take the yielding speed to be the maximum of the derivative of the instantaneous phase angle. Stronger crosslinks cause more catastrophic yielding and thus are represented by greater yielding speeds.

### Protection of cells during high shear-rate injection

Efficacy of directly-injected cell therapies is limited by the loss of viability of cells due to membrane disruption during injection^5^. Non-Newtonian injection media have been shown to improve the viability of injected cells due to protection from shear forces^6^, but design constraints that improve or worsen the viability for a given condition have not been established. It is currently hypothesized that the plug flow character of the gels during injection protects the cells from shear forces, or gradients in the flow velocity. Plug flow during the injection of DCHs may occur due to shear-banding or slip on the sides of the needle. We hypothesized that the character of the yielding transition would thus affect the viability of cells, as brittle yielding may introduce gradients in the velocity field which disrupt cell membranes. To assess the protection of cells from injection, we loaded HUVEC (Human umbilical vein epithelial cells) into each DCH material and injected them at clinically relevant flow rates through a 30G needle to impose a high shear-rate condition. We then measured the relative viability of the cells versus non-injected controls (**Figure 5, Supplemental Figure 7**). In accordance with our hypothesis, we found that the PNP hydrogel, which exhibits the most ductile yielding, affords the greatest protection of cells, at a level consistent with previous studies^6^. We found that more brittle yielding causes the cells to experience more membrane disruption due to velocity gradients and thus the Ca/Alginate and HA-ELP hydrogels caused lower cell viability upon injection. Different cell lines may have different levels of sensitivity to membrane disruption, so each cell type for a specific therapy could require a unique material selection for optimal use. These findings provide a design criterion for biomaterials which encapsulate cells for direct-injection therapies, as well as a verification of the importance of characterizing the yielding transition of dynamic hydrogels for translational applications.

**Figure 5.**
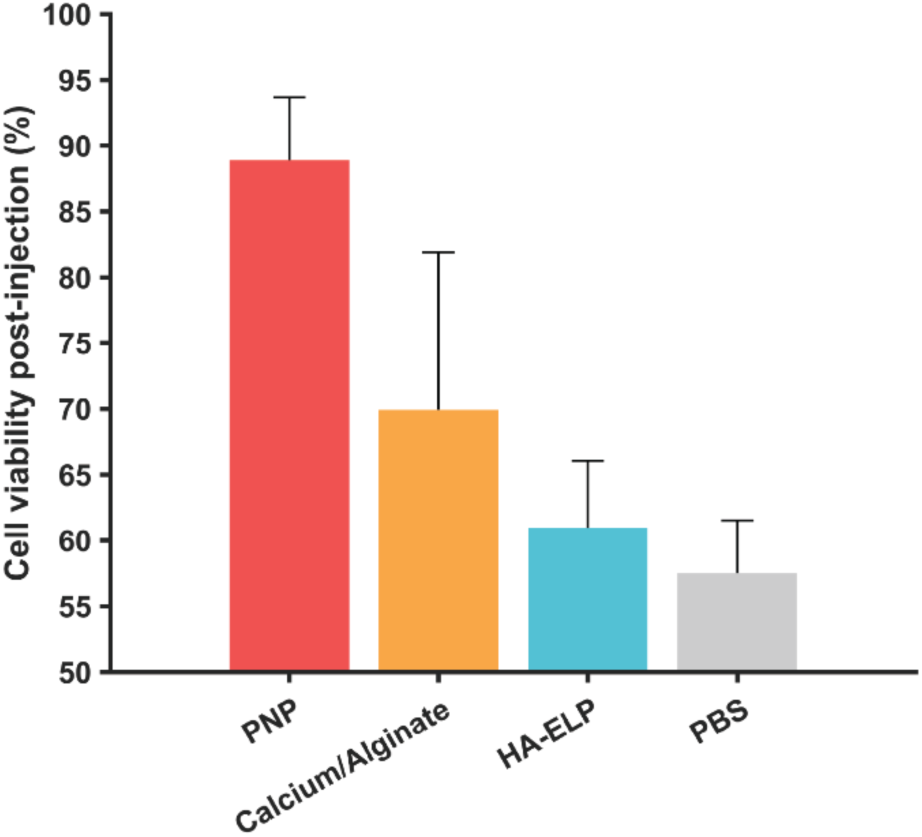
Viability of encapsulated HUVEC cells post high-shear rate injection. HUVEC cells were encapsulated at a concentration of 1.5 × 10^5^ cells/mL in each DCH as well as a PBS control, and injected at a clinically relevant flow rate (3 mL/min, shear rate ∼2.5 × 10^5^ s^-1^, Supplemental Section 1) through a 30 gauge needle. Cells were then stained with Calcein AM stain, and the viability of injected cells was compared against a non-injected control. Percent viability was calculated as the ratio of living cells between the injected and non-injected groups. Data is shown as mean of 5 measurements +/- SEM.

## Conclusion

Characterizing and understanding the yielding transition of soft materials has been a long-standing challenge. Sophisticated rheological techniques have recently been developed which attempt to model or apply greater understanding to the rheological response in the nonlinear regime, but these are often applied to model materials with relatively simple architectures. Moreover, less attention is given to the transition regime of yielding itself, especially the time-varying response of materials in the transition regime. Here, we applied fundamental rheological techniques to assess the yielding transition of three different dynamically associating hydrogel networks which have relevance for translational use in various biomedical applications. Through large-amplitude oscillatory shear experiments and subsequent Sequence of Physical Processes analysis, we were able to differentiate and quantify critical differences between smooth and catastrophic yielding in these dynamic hydrogels, dictated by the relative strength of the hydrogel crosslinks. We documented for the first time an elastically-derived stress overshoot in dynamic hydrogels using LAOS, corresponding to an elastic strain stiffening due to the nonlinearly increasing stiffness of polymer chain extension. We showed that the materials with smoother yielding transitions afforded greater protection to entrapped cells from shear forces or heterogeneities in the flow field introduced by fracture-like yielding. Most importantly, we intend this work to be a demonstration and accessible guide for of the application of sophisticated rheological techniques for investigators creating novel dynamic materials for cell encapsulation. Further work might assess the importance of the elastic stress overshoot in mechanotransduction of encapsulated cells, since these dynamically crosslinked materials are globally shear-thinning but may exhibit strain hardening on the length scale of an individual cell. Additionally, further work may seek to connect kinetic parameters, such as k_on_ and k_off_, of the individual crosslinks to the behavior of the yielding transition. Overall, this investigation highlights the importance of complete mechanical characterization of translational biomaterials.

## Experimental Procedures

### Resource availability

Please contact Prof. Eric Appel with any requests.

### Data and code availability

Computer code used to generate figures and process rheological data is freely and openly available online (https://github.com/neckman99/yielding_dynamic_hydrogels). All data reported in the paper will be shared by the corresponding author upon request.

### Preparation of polymer/nanoparticle hydrogels

Polymer/nanoparticle (PNP) hydrogels were synthesized by physical mixing of a hydrophobically-modified hydroxypropyl methylcellulose (HPMC) polymer (400 kDa) with core-shell (block)-polyethylene glycol-polylactic acid nanoparticles (NPs). Synthesis and characterization of this material has been reported previously^32^. After physical mixing in an Eppendorf tube, the gels were centrifuged to remove bubbles and left at 4C overnight.

### Preparation of calcium/alginate hydrogels

Ultrapure sodium alginate was purchased from NovaMatrix (PRONOVA UP LVG). The sodium alginate was dissolved at 4 wt% in phosphate buffered saline (PBS). Alginate stock solution was rapidly physically mixed with two syringes and an elbow mixer (McMaster-Carr) with a slurry of calcium sulfate powder in HBSS. Final gels were formulated at 1 wt% alginate polymer and 15 mM Ca^2+^. Rheological measurements were taken directly after the gel was observed to be homogeneous.

### Preparation of hyaluronic acid/elastin-like protein hydrogels

Hyaluronic acid (HA), modified with 12% aldehyde groups and with a molecular weight of 100 kDa, was dissolved in phosphate-buffered saline (PBS) at a concentration of 1 wt%. Similarly, elastin-like protein (ELP) modified with hydrazine was dissolved in PBS at the same concentration. Both solutions were stored at 4°C prior to use. Gels were formulated by physically mixing the HA and ELP solutions in an Eppendorf tube. The gel was centrifuged to remove air bubbles, then left overnight at 4C before rheological measurements.

### Rheometry

Rheological measurements were collected on a TA DHR-2 rheometer. All measurements were taken with a serrated 20mm parallel plate geometry to minimize slip artifacts, along with a water solvent trap to prevent evaporation. A gap size of 600 μm was used for all runs. To ensure the no slip condition was being attained, we frequency sweeps were run at two different gap sizes to ensure identical measurements. Frequency sweeps were run from 0.1Hz to 100 Hz at 0.5% amplitude for alginate gels and 1% amplitude for PNP and HA-ELP gels, in the linear viscoelastic regime for all materials. LAOS measurements were taken by running a logarithmic amplitude sweep from low to high strain amplitude while saving the instantaneous stress and strain. Amplitude sweeps were run at a frequency of 0.1 Hz to prevent inertial artifacts. After the run, the geometry was visually inspected to confirm that material had not been ejected from between the plates.

### Sequence of Physical Processes Analysis

Sequence of Physical Processes (SPP) analysis (Rogers *et al.* 2019)^22^ was carried out on the time-varying stress and strain data exported from the rheometer, using MATLAB software. The stress curves were fit using Fourier analysis to n=5 odd harmonics, which was qualitatively observed to be the appropriate number of harmonics which fit the data without overfitting noise or introducing numerical instability. The software produced time-varying modulus and phase angle data. The odd Fourier fit to the stress curve was used for subsequent stress decomposition analysis.

### Cell viability after high-strain rate injection

Human umbilical vein epithelial cells (HUVEC, ATCC) were cultured in GM-2 Endothelial Cell Growth Medium (Lonza) supplemented with 1% Pen-Strep at 37° C and 5% CO_2_. Cells were passaged at 80% confluence and all passage numbers used in this study were below 10. For encapsulation, cells were trypsinized with 0.05% Trypsin/EDTA, centrifuged at 300g for 5 minutes, and then counted and resuspended in PBS to an appropriate volume. For each gel, a cell/PBS slurry was added to the liquid precursor followed by gentle physical mixing of the gels. The gels were allowed to rest for one hour on ice before injection. 100 μL of gel/cell mixture was injected using a syringe pump at 3.0 mL/min through a 30G needle (I.D. = 0.159 mm) in triplicate onto a 24-well plate pre-loaded with a solution of Calcein-AM at 10 μM (Santa Cruz Biotechnology). The plates were centrifuged at 50g for 1 minute to ensure good surface contact for microscope imaging. After 1 hour incubation at 37 °C confocal microscope images (Leica, 10x objective) were taken of the entire gel surfaces in one representative focal plane.

## Author Contributions

N.E., A.K.G., and E.A.A., conceptualization; N.E., A.K.G., G.J., M.S.H., K.K., methodology; N.E., A.K.G., and K.K., data analysis; N.E., writing; E.A.A. and S.R., paper review and editing; E.A.A., S.R., S.C.H., supervision.

## Acknowledgements

N.E. was supported by a National Science Foundation GRFP, no. DGE-2146755. This work was supported in part by the Bill & Melinda Gates Foundation (INV-027411). The authors thank Dr. Sarah Hull for providing HA-ELP materials and advice on gel preparation. The authors thank Dr. Alex Prossnitz, Dr. Sophia Bailey, and Dr. Julie Baillet for their helpful discussions and advice.

## References

1. Webber, M.J., Appel, E.A., Meijer, E.W., and Langer, R. (2016). Supramolecular biomaterials. Nature Mater 15, 13–26. 10.1038/nmat4474.

2. Correa, S., Grosskopf, A.K., Lopez Hernandez, H., Chan, D., Yu, A.C., Stapleton, L.M., and Appel, E.A. (2021). Translational applications of hydrogels. Chemical Reviews 121, 11385– 11457.

3. Zhang, K., Feng, Q., Fang, Z., Gu, L., and Bian, L. (2021). Structurally Dynamic Hydrogels for Biomedical Applications: Pursuing a Fine Balance between Macroscopic Stability and Microscopic Dynamics. Chem. Rev. 121, 11149–11193. 10.1021/acs.chemrev.1c00071.

4. Sisso, A.M., Boit, M.O., and DeForest, C.A. (2020). Self-healing injectable gelatin hydrogels for localized therapeutic cell delivery. Journal of Biomedical Materials Research Part A 108, 1112–1121. 10.1002/jbm.a.36886.

5. Aguado, B.A., Mulyasasmita, W., Su, J., Lampe, K.J., and Heilshorn, S.C. (2012). Improving Viability of Stem Cells During Syringe Needle Flow Through the Design of Hydrogel Cell Carriers. Tissue Engineering Part A 18, 806–815. 10.1089/ten.tea.2011.0391.

6. Lopez Hernandez, H., Grosskopf, A.K., Stapleton, L.M., Agmon, G., and Appel, E.A. (2019). Non-Newtonian Polymer–Nanoparticle Hydrogels Enhance Cell Viability during Injection. Macromolecular Bioscience 19, 1800275. 10.1002/mabi.201800275.

7. Morley, C.D., Ding, E.A., Carvalho, E.M., and Kumar, S. (2023). A Balance between Inter- and Intra-Microgel Mechanics Governs Stem Cell Viability in Injectable Dynamic Granular Hydrogels. Advanced Materials 35, 2304212. 10.1002/adma.202304212.

8. Tanaka, F., and Edwards, S.F. (1992). Viscoelastic properties of physically crosslinked networks: Part 2. Dynamic mechanical moduli. Journal of Non-Newtonian Fluid Mechanics 43, 273–288. 10.1016/0377-0257(92)80028-V.

9. Yu, A.C., Lian, H., Kong, X., Lopez Hernandez, H., Qin, J., and Appel, E.A. (2021). Physical networks from entropy-driven non-covalent interactions. Nat Commun 12, 746. 10.1038/s41467-021-21024-7.

10. Marco-Dufort, B., Iten, R., and Tibbitt, M.W. (2020). Linking Molecular Behavior to Macroscopic Properties in Ideal Dynamic Covalent Networks. Journal of the American Chemical Society 142, 15371–15385. 10.1021/JACS.0C06192/ASSET/IMAGES/LARGE/JA0C06192_0009.JPEG.

11. Nicolella, P., and Seiffert, S. (2022). Mechanical switching of a comblike dual dynamic polymer network. Journal of Rheology 66, 1153–1161. 10.1122/8.0000388.

12. Song, J., Li, Q., Chen, P., Keshavarz, B., Chapman, B.S., Tracy, J.B., McKinley, G.H., and Holten-Andersen, N. (2022). Dynamics of dual-junction-functionality associative polymer networks with ion and nanoparticle metal-coordinate cross-link junctions. Journal of Rheology 66, 1333–1345. 10.1122/8.0000410.

13. Grosskopf, A.K., Saouaf, O.A., Hernandez, H.L., and Appel, E.A. (2021). Gelation and yielding behavior of polymer–nanoparticle hydrogels. Journal of Polymer Science 59, 2854– 2866. 10.1002/POL.20210652.

14. Herbst, F., Döhler, D., Michael, P., and Binder, W.H. (2013). Self-Healing Polymers via Supramolecular Forces. Macromolecular Rapid Communications 34, 203–220. 10.1002/marc.201200675.

15. Yasui, T., Zheng, Y., Nakajima, T., Kamio, E., Matsuyama, H., and Gong, J.P. (2022). Rate-Independent Self-Healing Double Network Hydrogels Using a Thixotropic Sacrificial Network. Macromolecules 55, 9547–9557. 10.1021/acs.macromol.2c01425.

16. Ollier, R.C., and Webber, M.J. (2024). Strain-Stiffening Mechanoresponse in Dynamic-Covalent Cellulose Hydrogels. Biomacromolecules. 10.1021/acs.biomac.4c00450.

17. Griebler, J.J., Donley, G.J., Wisniewski, V., and Rogers, S.A. (2024). Strain shift measured from stress-controlled oscillatory shear: Evidence for a continuous yielding transition and new techniques to determine recovery rheology measures. Journal of Rheology 68, 301–315. 10.1122/8.0000756.

18. Park, J.D., and Rogers, S.A. (2018). The transient behavior of soft glassy materials far from equilibrium. Journal of Rheology 62, 869–888. 10.1122/1.5024701.

19. Klein, C.O., Spiess, H.W., Calin, A., Balan, C., and Wilhelm, M. (2007). Separation of the Nonlinear Oscillatory Response into a Superposition of Linear, Strain Hardening, Strain Softening, and Wall Slip Response. Macromolecules 40, 4250–4259. 10.1021/ma062441u.

20. Graham, M.D. (1995). Wall slip and the nonlinear dynamics of large amplitude oscillatory shear flows. Journal of Rheology 39, 697–712. 10.1122/1.550652.

21. Rogers, S.A. (2017). In search of physical meaning: defining transient parameters for nonlinear viscoelasticity. Rheol Acta 56, 501–525. 10.1007/s00397-017-1008-1.

22. Donley, G.J., Bruyn, J.R. de, McKinley, G.H., and Rogers, S.A. (2019). Time-resolved dynamics of the yielding transition in soft materials. Journal of Non-Newtonian Fluid Mechanics 264, 117–134. 10.1016/J.JNNFM.2018.10.003.

23. Kamani, K.M., and Rogers, S.A. (2024). Brittle and ductile yielding in soft materials. Proceedings of the National Academy of Sciences 121, e2401409121. 10.1073/pnas.2401409121.

24. Zhang, V., Kang, B., Accardo, J.V., and Kalow, J.A. (2022). Structure-Reactivity-Property Relationships in Covalent Adaptable Networks. J Am Chem Soc 144, 22358–22377. 10.1021/jacs.2c08104.

25. Sikorski, P., Mo, F., Skjåk-Bræk, G., and Stokke, B.T. (2007). Evidence for Egg-Box-Compatible Interactions in Calcium−Alginate Gels from Fiber X-ray Diffraction. Biomacromolecules 8, 2098–2103. 10.1021/bm0701503.

26. Roth, J.G., Huang, M.S., Navarro, R.S., Akram, J.T., LeSavage, B.L., and Heilshorn, S.C. (2023). Tunable hydrogel viscoelasticity modulates human neural maturation. Science Advances 9, eadh8313. 10.1126/sciadv.adh8313.

27. Kölmel, D.K., and Kool, E.T. (2017). Oximes and Hydrazones in Bioconjugation: Mechanism and Catalysis. Chem Rev 117, 10358–10376. 10.1021/acs.chemrev.7b00090.

28. Olsen, B.D., Kornfield, J.A., and Tirrell, D.A. (2010). Yielding Behavior in Injectable Hydrogels from Telechelic Proteins. Macromolecules 43, 9094–9099. 10.1021/ma101434a.

29. Kwang Soo Cho, Cho, K.S., Kyu Hyun, Hyun, K., Kyung Hyun Ahn, Ahn, K.H., Jeong, Y., Seung Jong Lee, and Lee, S.J. (2005). A geometrical interpretation of large amplitude oscillatory shear response. Journal of Rheology 49, 747–758. 10.1122/1.1895801.

30. Rogers, S.A., and Lettinga, M.P. (2012). A sequence of physical processes determined and quantified in large-amplitude oscillatory shear (LAOS): Application to theoretical nonlinear models. Journal of Rheology 56, 1–25. 10.1122/1.3662962.

31. Donley, G.J., Singh, P.K., Shetty, A., and Rogers, S.A. (2020). Elucidating the G″ overshoot in soft materials with a yield transition via a time-resolved experimental strain decomposition. Proceedings of the National Academy of Sciences 117, 21945–21952. 10.1073/pnas.2003869117.

32. Grosskopf, A.K., Roth, G.A., Smith, A.A.A., Gale, E.C., Hernandez, H.L., and Appel, E.A. (2020). Injectable supramolecular polymer–nanoparticle hydrogels enhance human mesenchymal stem cell delivery. Bioengineering & Translational Medicine 5, e10147. 10.1002/btm2.10147.

