## Supplementary Materials for "Crosslink Strength Governs Yielding Behavior in Dynamically Crosslinked Hydrogels"

### Dynamic Hydrogel Crosslink Strength Controls Yielding Characteristics and Cell Viability During Injection


### Table of Contents


Calculation of network crosslink densities……………………………………………………….……..2
Calculation of apparent strain rate during injection…………………………………………….……2
Supplemental Figure 1. Flow sweeps of dynamic hydrogel materials………………………………3
Supplemental Figure 2. Stress/strain Lissajous curves from LAOS experiments……………….…4
Supplemental Figure 3. LAOS experiment for Carbopol 980…………………………………………5
Supplemental Figure 4. Alternate view of the 3D LAOS curves of HA-ELP………………………..5
Supplemental Figure 5. Raw stress/time data from the flow regime in LAOS experiments……...6
Supplemental Figure 6. Elastic strain stiffening calculation illustration…………………………..…6
Supplemental Figure 7. Cell viability upon injection in injectable hydrogels…………………………7
Code Availability……………………………………………………………………….............…………....7
Supplemental References…………………………………………………………………………….…...8

**Calculation of network crosslink densities.**

For the calculation of the crosslink density of the Polymer/Nanoparticle gel, we assume that one binding pair consists of a persistence length of HPMC-C12 (approximately 10 nm) and a surface of nanoparticle of 100 $nm^2$. Since $K_{eq}$>>1, we assume all binding sites are occupied at equilibrium. The NP diameter is 30 nm[1], but we assume the polymer binds to the PLA core, which has a radius of 20 nm accounting for the length of the PEG chains. The number of binding sites per mL is thus

$$\text{NP binding site concentration} = \text{(NP mass concentration)} * \frac{m_{PLA}}{m_{NP}} * \rho_{PLA} * \frac{1}{V_{core}} * 4\pi \left(\frac{d_{core}}{2}\right)^2$$

Which yields a molarity of 1.6 mM for a 5% NP gel by mass. To calculate the number of polymer binding sites per mL, we take the persistence length of 10 nm to be one binding site, made up of 10 monomers of average molecular weight 191.87 g/mol. The calculation is

$$\begin{aligned}&\text{Polymer binding site concentration}\\ &\quad = \text{(polymer mass concentration)}\\ &\quad * \frac{1}{\text{(monomer MW)} * \text{(\# monomers per persistence length)}}\end{aligned}$$

Which yields a molarity of 5.2 mM, which is in excess of the NP binding site molarity, so the gel crosslink molarity is approximately 1.6 mM.

For the calculation of Ca/Alginate crosslink density, we assume all Ca ions are involved in crosslinking, and that each crosslink “egg-box” occupies 4 $Ca^{2+}$ ions. The density is then 15 mM/4 ≈ 3.8 mM.

For the HA-ELP gels, the HA is modified with 10% hydrazine on a monomer basis, and we assume all groups form crosslinks at equilibrium, yielding a crosslink density of

$$\text{Polymer binding site concentration} = \text{(polymer mass concentration)} * \frac{\text{\% modification}}{\text{(HA monomer MW)}}$$

Which gives a crosslink density of 2.6 mM.

**Calculation of apparent strain rate during injection.**

To calculate the maximum strain rate experienced by the gels during injection, we calculate the apparent strain rate according to the geometry and flow rate and then apply the Rabinowitch correction:

$$\dot{\gamma} = \dot{\gamma}_{app}\left(\frac{3n+1}{4n}\right) = \frac{4Q}{\pi R^3}\left(\frac{3n+1}{4n}\right) \approx 2.5 * 10^5\ s^{-1}$$

Where Q = 3 mL/min, n ~ 0.2 (Supplemental Figure 1), and R = 0.15mm.

**Supplemental Figures.**

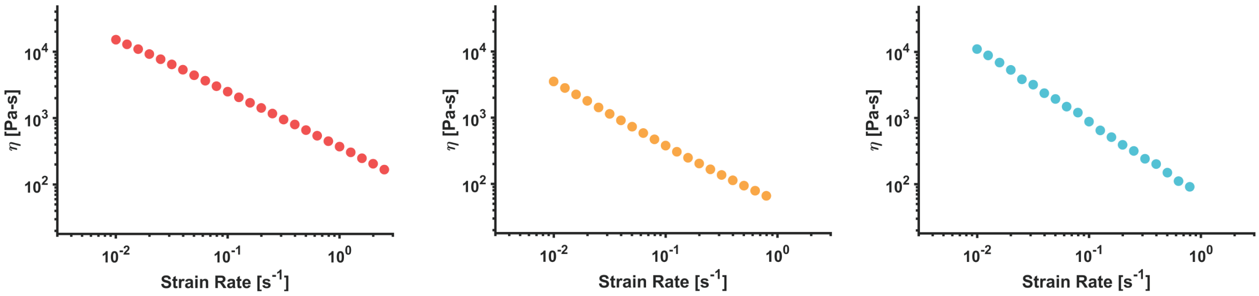


**Figure S8.** Steady-state flow sweeps show the shear thinning behavior of all DCH materials (Polymer/Nanoparticle, left; Ca/Alginate, middle; HA-ELP, right).

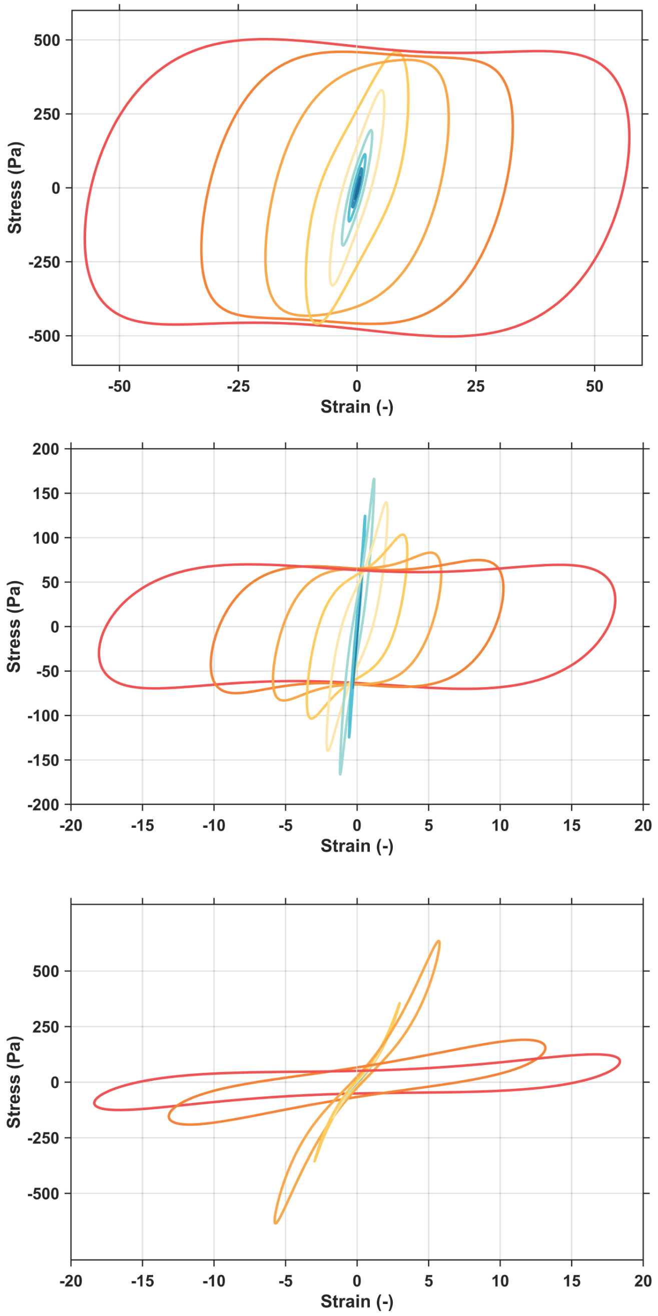


**Figure S9.** Stress/strain Lissajous curves from LAOS experiments for PNP (top), Ca/Alginate (middle), and HA-ELP (bottom).

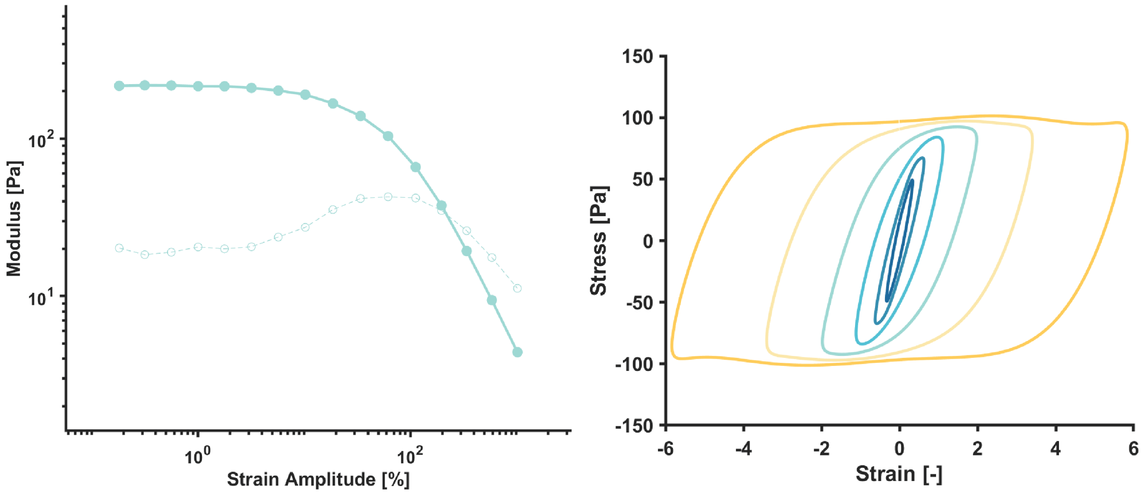


**Figure S10.** LAOS experiment for Carbopol 980 (20 wt%). The amplitude sweep shows a G'' overshoot but no corresponding stress overshoot in the Lissajous curves.

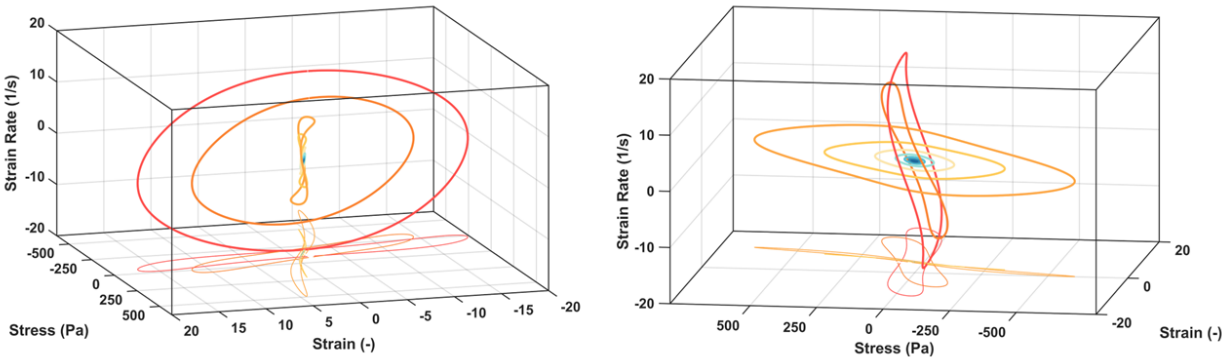


**Figure S11.** Alternate view of the 3D LAOS curves of HA-ELP to show the yielding transition between orthogonal planes.

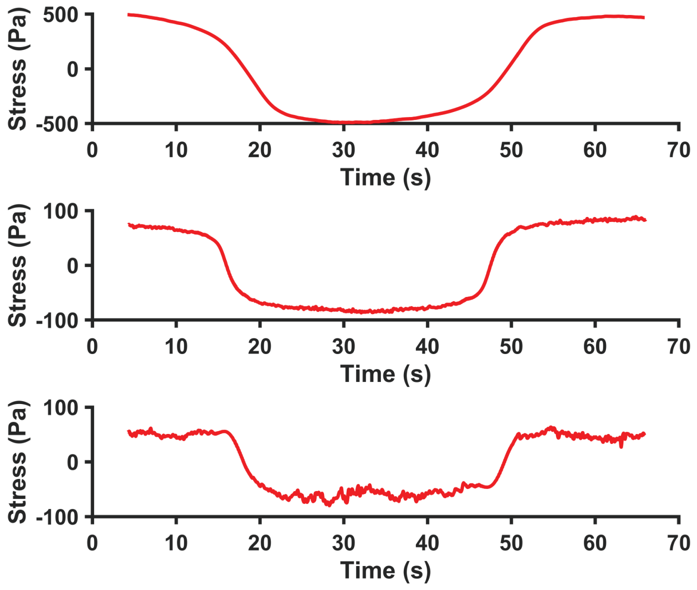


**Figure S12.** Raw stress/time data from the flow regime in the LAOS experiment for Polymer/Nanoparticle gel (top), Ca/Alginate gel (middle), and HA-ELP gel (bottom). Flattening of the stress curves shows the shear-banding response at post-yield conditions.

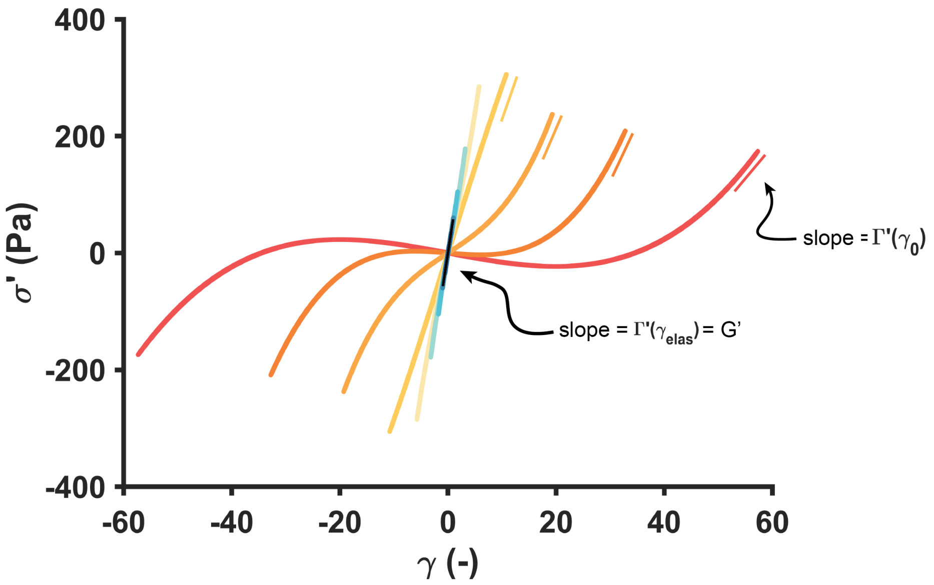


**Figure S13.** Illustration of the calculation of main text Figure 3(c), with representative data from the polymer/nanoparticle hydrogel.

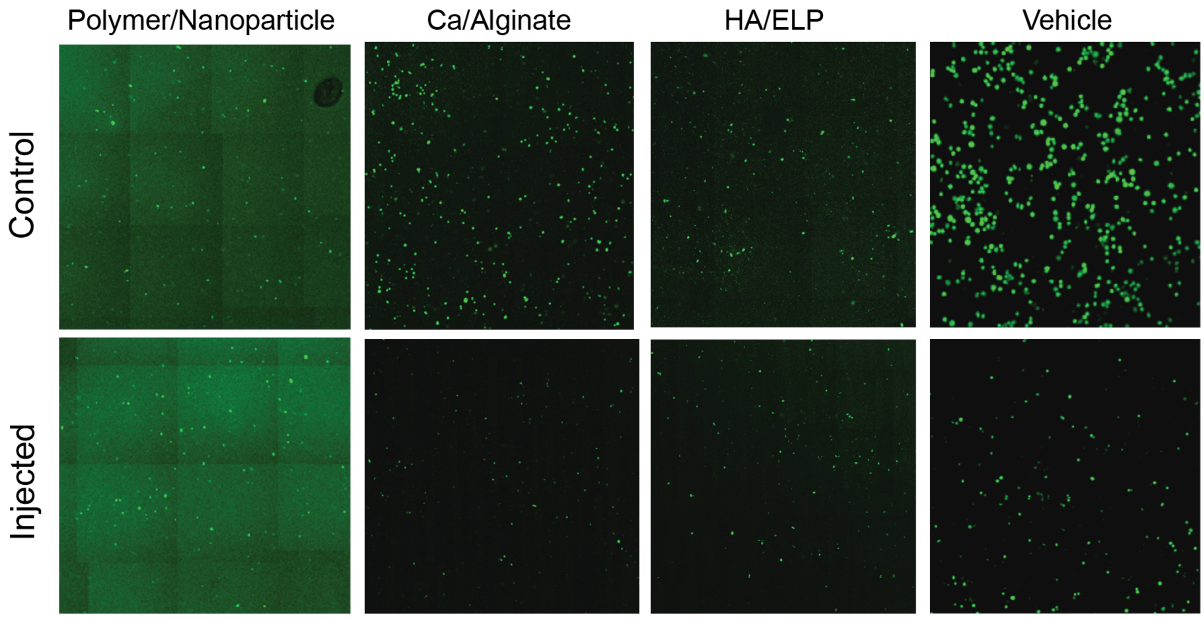


**Figure S14.** Representative confocal images with Calcein AM (live cell) stain for injected gels and PBS control. Viabilities were calculated by relative counts between the same material.

**Code Availability.**

Code used to generate figures and process rheological data is openly and freely available at https://github.com/neckman99/yielding_dynamic_hydrogels.

**Supplemental References.**


1. Grosskopf, A.K., Saouaf, O.A., Hernandez, H.L., and Appel, E.A. (2021). Gelation and yielding behavior of polymer–nanoparticle hydrogels. Journal of Polymer Science *59*, 2854–2866. https://doi.org/10.1002/POL.20210652.
